# DTMol: Pocket-based Molecular Docking using Diffusion Transformers

**DOI:** 10.1101/2025.04.13.648103

**Authors:** Haotian Teng, Ran Wang, Yihang Shen, Ye Yuan, Carl Kingsford

## Abstract

In computational chemistry, molecular docking—predicting the binding structure of a small molecule ligand to a protein—is vital for understanding interactions between small molecules and their protein targets, with broad applications in drug discovery (Morris and Lim-Wilby, 2008). Traditional docking methods rely on energy-based scoring functions and optimization algorithms to identify ligand-binding structures. However, these methods are often slow and inaccurate due to the extensive search space for binding structures and the complexity of scoring function landscapes (Corso et al., 2023).

Deep learning techniques have emerged as promising alternatives for molecular docking. These methods harness the power of neural networks to understand the complex interactions between protein pockets and ligands from large datasets. They can be classified into two categories: regression-based methods (Stark et al., 2022; Lu et al., 2022), which offer greater computational efficiency compared to traditional docking methods but have yet to offer substantial improvements in accuracy (Corso et al., 2023); and generative models, particularly diffusion models (Corso et al., 2023; Qiao et al., 2024; Nakata et al., 2023; Lu et al., 2024; Schneuing et al., 2024), which lead to significant accuracy improvements compared to regression-based methods.

Most diffusion-based docking methods aim to address the blind docking problem, where the ligand is docked without prior knowledge of the specific protein pocket—a site of the protein with potential ligand-binding capabilities—and only the overall protein structure is provided (Yim et al., 2024). However, in many drug discovery cases, specific pockets have already been identified (Zheng et al., 2013), meaning that the docking process only needs to determine how the ligand binds to the pocket. This defines the pocket-based docking problem. Pocket-based docking leverages the physical and chemical information of protein side chains around the pocket, significantly reducing the search space for ligand binding structures and improving the understanding of atomic-level interactions. Additionally, focusing on the local protein substructure rather than the global structure enhances computational efficiency, which allows for the use of more complex neural architectures.

We introduce DTMol, a novel diffusion model designed to tackle the pocket-based docking problem. Our model integrates a pretrained molecular representation framework with a diffusion transformer architecture. The advantages of this design are twofold: first, the molecular representation models we employ are pretrained on separate datasets of small molecules and protein pockets, which are larger than the datasets of ligand-pocket interactions. Consequently, these pretrained models encode typical structural features of both elements, enhancing our model’s ability to represent them and predict binding structures. Second, the transformer architecture used in our diffusion model is more capable of effectively capturing all-atom interactions between the pocket and the ligand.

We test our method on the docking benchmark PDBBind v2020 and PoseBuster, and compare it with other methods. DTMol achieves 77.65% top-1 success rate (RMSD ¡ 2Å) on PoseBuster, outperforms the second-best docking software Gnina (65.65%) We also test the performance of our method on a real-world virtual screening task using Janus kinase 2 (PDB ID: 6BBV) as the target protein. A TR-FRET screening experiment is performed on the union of top ligands identified via physics-based docking and the diffusion model, in order to measure their inhibitory activity against the JAK2 protein. The results shows that our model achieved the only positive rank correlation score compared to other traditional docking methods and machine learning-based methods.

To summarize, the main contributions of this work are:

- We introduce DTMol, a novel diffusion model designed to address the pocket-based docking problem. This model represents the first application of the diffusion transformer architecture in solving pocket-based molecular docking.
- We propose a novel SE(3)-equivariant transformer architecture with an SE(3)-invariant and a parallel SE(3)-equivariant flow, which is straightforward to implement and train while achieving excellent performance.
- We achieve new state-of-the-art results on both the PDBBind and PoseBuster pocket-based docking benchmarks, obtaining top-1 prediction rates of 45.05% and 77.65%, respectively, at RMSD *<* 2Å.
- Doing virtual screening on Janus kinase 2, DTMol is able to achieve the only positive rank score; and identifies 2 new experimentally effective JAK2 inhibitors.

## 1 Background

Diffusion models are a class of generative models that learn to iteratively denoise a signal, starting from random noise and gradually refining it into a meaningful output [10, 11]. The noising process is defined by a Markov process of diffusion steps, where each step adds a small amount of noise to the signal. The reverse process, used for generating samples, is learned by training a neural network to predict the noise added at each step, conditioned on the noisy signal.

The forward noising process of diffusion model can be seen as stochastic differential equation (SDE)

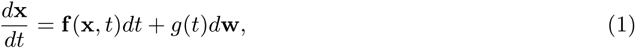

which gradually noises an input data **x**(*t* = 0) ∼ *p*(**x**) from data distribution *p*(**x**), where **w** is the wiener process (Song et al., 2021). Different choices of the draft term **f** (*x, t*) and noise term *g*(*t*) would result in different forms of diffusion framework; this includes *Score matching with Langevin dynamics* (SMLD), where **f** (**x**, *t*) = 0 and 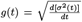 (Song and Ermon, 2019), and *Denoising diffusion probabilistic modeling* (DDPM), with 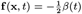 and 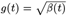 (Sohl-Dickstein et al., 2015; Ho et al., 2020). The generative process is regarded as solving the reverse SDE

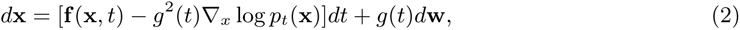

where *p*_*t*_(**x**) is the probability distribution of noised input **x**(*t*). It is generally intractable to directly compute the score ∇_*x*_ log *p*_*t*_(**x**) and this is usually approximated by a neural network **s**_*θ*_(**x**, *t*) (Song et al., 2021), which is known as score matching (Vincent, 2011)

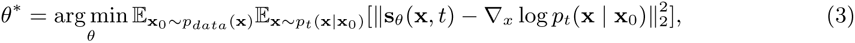

where the hard-to-calculate marginal distribution *p*_*t*_(**x**) is approximated by a conditional distribution on sampled input **x**_0_. It is beneficial to give the time discrete form of the forward process, where *x*_*t*_ is the noisy signal at step *t, x*_0_ is the original signal, and **z**_*t*_ is the noise added at step *t*. For SMLD, 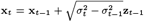, for DDPM we have 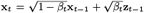, where **z** ∼ 𝒩 (0, **I**).

We can have the posterior form of distribution *p*(**x**_*t*_ | **x**_0_) for any given *t* from the recurrence form *p*(**x**_*t*_ | **x**_*t*−1_) (see detailed derivations in Appendix), for SMLD *P* (**x**_*t*_|**x**_0_) = 𝒩 (**x**_*t*_; **x**_0_, *σ*_*t*_**I**) and DDPM 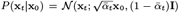, where 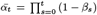. Note that for DDPM, if the input **x**_0_ is already scaled to have a unit norm, then **x**_*t*_ at any time *t* will also maintain a unit norm. Therefore, DDPM is referred to as a variance-preserved diffusion process, in contrast to SMLD where the variance of **x** increases over time. Both approaches can be useful, depending on the desired behavior of the diffusion process. In our case, we use the DDPM form to add perturbation noise to the atom coordinates because we want to preserve the variance of atom distribution during the diffusion process. We use the SMLD form for adding noise to the rotation Euler vector and translation vector, as these vectors are initially zero.

### Traditional score-based docking methods

Traditional search-based docking methods (Trott and Olson, 2010; Halgren et al., 2004; Thomsen and Christensen, 2006) aim to find the optimal binding pose of a ligand within a protein pocket by exploring the conformational space. These methods typically consist of a scoring function to evaluate the quality of a ligand pose and a search algorithm to explore the conformational space. Scoring functions can be force field-based, empirical, or knowledge-based, considering factors such as van der Waals interactions, electrostatic interactions, hydrogen bonding, and hydrophobic interactions. While traditional search-based docking methods have been widely used and have shown success in drug discovery, they often struggle with the high dimensionality and complexity of the conformational space, leading to limitations in computational efficiency and accuracy (Stärk et al., 2022). Additionally, the scoring functions may not always capture the full complexity of ligand-protein interactions, potentially resulting in inaccuracies in pose prediction (Zhou et al., 2022).

### Machine learning for molecular docking

Recently machine learning methods have been applied to solve the molecular docking problem. DeepDock (Méndez-Lucio et al., 2021) substitutes score functions used in traditional docking methods with a learned potential, which, when combined with optimization algorithms, generates binding conformations of ligands. EquiBand (Stärk et al., 2022) and TANKBind (Lu et al., 2022) use regression-based techniques to predict the docking poses of ligands. As noted by Corso et al. (2023), these regression-style methods that generate only a single pose per ligand tend to produce the mean of many viable alternative poses, increasing the likelihood of yielding physically implausible poses. Uni-Mol (Zhou et al., 2022) uses the transformer architecture to tackle molecular representation learning (MRL), which can also be adapted for the docking problem through fine-tuning. DiffDock (Corso et al., 2023) is an approach closely related to our method. However, Diff-Dock is not a pocket-based approach and neglects protein side-chain conformations, while our method is pocket-based and models side chains.

### Diffusion Transformers

Transformer architectures have shown great success in various sequence-to-sequence tasks, including machine translation and language modeling. Transformers are well-suited for generation tasks by training using simple next-token prediction. Recent work has integrated transformers with diffusion models, known as diffusion transformers (Peebles and Xie, 2023). Diffusion transformers have been applied to image synthesis (Rombach et al., 2022) and video generation (Brooks et al., 2024). In the context of molecular modeling, several diffusion generative models have been developed and applied to several tasks, including molecular generation (Hoogeboom et al., 2022), conformer generation (Xu et al., 2021; Jing et al., 2022), protein design (Trippe et al., 2022) and molecular docking (Corso et al., 2023). However, all these models used a *SE*(3)-equivalent Graph Neural Network (GNN), potentially limiting the model performance. The transformer can be viewed as a kind of more general GNN, where the underlying graph is complete and an attention mechanism is used as the node surrogation strategy. The transformer architecture can learn to attend to different parts of the protein pocket and the generated molecule, enabling the model to capture the spatial and chemical constraints governing protein-ligand interactions. In addition, the architecture of the transformer needs to be redesigned to make the model *SE*(3)-equivariant which is required for molecular docking tasks (See the Model Detail section below).

## 2 Molecular Docking with Diffusion Transformer

Recent work such as DiffDock (Corso et al., 2023) has treated molecular docking as a generative modeling problem, where molecular conformation is generated given the protein. While DiffDock has shown strong improvements in docking accuracy, we argue that the generative model has not yet reached its full potential in molecular docking tasks. DiffDock generates molecule conformations at any potential binding site across the entire protein, achieving blind molecular docking. Although this approach has some benefits, it requires the model to take the whole protein as input, which could constrain the choice of neural network architectures that can be used, since the number of atoms of whole protein could be potentially large. DiffDock addresses this constraint by using only the *α*C of the protein backbone and employing a simple *SE* (3) convolutional neural network. We argue that a whole protein input for molecular docking may not be necessary, as the protein binding site can be detected beforehand, allowing for molecular docking at each binding site separately. Thus, we choose to include only the protein pocket as input, significantly reducing the number of atoms that need to be considered. This simplification allows us to use a more sophisticated transformer score network and consider all the atoms except hydrogen of the given protein pocket.

### 2.1 Overview

We regard the molecule as a point cloud where each point is an atom of the molecule. A ligand or a protein pocket of *n* atoms represented by *n* coordinates in ℝ^3^ results in an element in coordinate space 𝒳 = ℝ. Translation of ligand positions is defined as an element in the 3D translation group 𝕋 (3), and the rigid rotations of ligand is defined as an element in the 3D rotation group *SO*(3). Instead of using torsional rotations as DiffDock, we used an atom-wise perturbation, adding Gaussian noise to each atom during the diffusion process.

Specifically, given a ligand with the position **x** ∈ ℝ^3*n*^, an operation *α* is used to represent either the translation operation *α*^*tr*^ or a rotation operation *α*^*rot*^. A translation action *α*^*tr*^ : 𝕋 (3) × ℝ^3*n*^ → ℝ^3*n*^ is defined as *α*^*tr*^(**r, x**)_*i*_ = **x**_*i*_ + **r**, where **x**_*i*_ ∈ ℝ^3^ is the coordinate of the *i*-th atom and **r** ∈ 𝕋 (3) ≅ ℝ^3^ is a translation vector. It means the molecule would be translated uniformly by **r**. The rotation operation *α*^*rot*^ : *SO*(3) × ℝ^3*n*^ → ℝ^3*n*^ is defined as 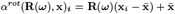, where ***ω*** ∈ *SO*(3) is an element in the rotation group, **R**(***ω***) is the corresponding rotation matrix, and 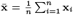 is the center of mass of the ligand. Finally, for the atom-wise perturbation noise, we sample *n* independent noises ***ϵ***_*i*_ from zero-mean Gaussian distribution in ℝ^3^, and add these noises to each atom, leading to 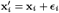 for each atom.

Combined together, at time step *t*, the diffusion process includes sampling a translation vector **r**_*t*_ ∼ 𝒩 (0, *σ*_*tr*_ (*t*)^2^ **I**), a rotation vector ***ω***_*t*_ ∼ *p*(*σ*_*rot*_(*t*)), and *n* perturbation noise vectors ***ϵ***_*i*_ ∼ 𝒩 (0, *σ*_*per*_ (*t*)^2^ **I**), and then applied to the pose of the ligand **x**. The sampling distribution *p*(*σ*_*rot*_(*t*)) for rotation space *SO*(3) is given in the next section.

### 2.2 Diffusion on *SO*(3) space

In our diffusion process, we model the rigid rotations of ligands as elements in the 3D rotation group *SO*(3). The diffusion kernel on *SO*(3) is given by the isotropic Gaussian distribution on the *SO*(3) group (*IGSO*(3)) (Nikolayev and Savyolov, 1997; Leach et al., 2022), which can be efficiently sampled using the axis-angle parameterization.

At each diffusion step *t*, we sample a rotation vector ***ω***_*t*_ ∈ ℝ^3^ from the *IGSO*(3) distribution, denoted as *p*(***ω***, *σ*_*rot*_(*t*)). The rotation vector ***ω***_*t*_ is represented by its axis-angle representation, ***ω***_*t*_ = *ω*_*t*_**ê**_*t*_, where *ω*_*t*_ ∈ [0, *π*] is the rotation angle and **ê**_*t*_ ∈ ℝ^3^ is a unit vector representing the rotation axis. The unit vector **ê**_*t*_ is sampled uniformly from the unit sphere in ℝ^3^, and the rotation angle *ω*_*t*_ is sampled according to the probability density function:

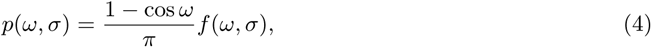

where *f* (*ω, σ*) is given by:

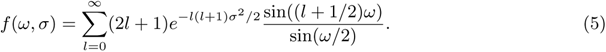

Further, the score of the diffusion kernel is ∇ ln 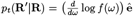, where **R**^′^ = **R**_***ω***_ **R** is theresult of applying the Euler vector ***ω*** = *ω***ê** to **R** (which is essentially the identity matrix at the beggining **R**(*t* = 0) = **I**) (Yim et al., 2023). The score computation and sampling is accomplished by precomputing the truncated infinite (*l* up to 5000) series and interpolating the CDF of *p*(*ω*), following Corso et al. (2023).

### 2.3 Perturbation noise

Instead of using the torsion diffusion process presented in previous work (Jing et al., 2022; Corso et al., 2023), we worked in the coordinate space with a perturbation diffusion process for several benefits. Working purely in the operation space, like DiffDock (Corso et al., 2023), makes the model sensitive to the seed conformation of the molecule. It would require a similar seed conformation for a molecule during training and inference for the torsion operator to have consistent meaning, which is usually not possible as the true conformation of a ligand binding to a protein is regarded as unknown when performing molecular docking. In DiffDock, this issue is addressed by generating an initial conformation using RDKit during inference. During the training phase, instead of using the true conformation of the ligand, a ligand structure is generated by RDKit and then aligned with the true conformation using a conformer matching procedure (Jing et al., 2022). This aligned conformation could introduce bias during training as it could differs from the true conformation. Using perturbation noise, on the other hand, avoids this issue, as it directly works on the coordinate space.

Diffusion in the operator space requires the operator to be a group action, which is true for translation and rotation, but not for torsional operations. This is because torsional changes break associativity, meaning that a molecule would have a different pose when applying the sequential operations individually compared to applying the composed operation *α*(*g*_1_*g*_2_, **x**) ≠ ss*α*(*g*_1_, *α*(*g*_2_, **x**)) (Corso et al., 2023).

This issue was ignored in DiffDock, as they approximate the torsional change to be a group action if the torsional rotation angle is small, while this isn’t always the case. On the other hand, perturbation noise, which adds zero-mean Gaussian noise to each atom, does not change the expected mean coordinates, thus keeping the translation and rotation operation group actions. (See the Appendix Proof section for a detailed derivation).

The torsion operation reduces the degrees of freedom of molecule conformation, which could be beneficial, as chemical bonds such as covalent bonds are generally energetically restricted, and the bond length usually does not change significantly (Axelrod and Gómez-Bombarelli, 2022; Jing et al., 2022). A full perturbation on all atoms would potentially introduce unrealistic changes that could alter the bond length or other fixed structures. However, we rarely observe that the perturbation noise results in unrealistic structures; we believe this is because of the use of a pretrained model on molecules, which has learned a good prior of general chemical bond knowledge. Structures that have not deviated significantly from realistic configurations can be corrected by an energy-based relaxation afterward. Although the covalent bond does not change much, it can still vary under different circumstances, which would not be reflected in a torsion-based diffusion method.

Despite the potential benefits of torsion operations in maintaining realistic molecular structures, they introduce additional complexity for score prediction, as torsion operations lie in a high-dimensional Tm Riemannian space, whereas perturbation operations are in the plain Euclidean space. Perturbation operations for each atom are independent of each other, while torsion operations are typically entangled with each other (Corso et al., 2023); a rotation in one bond would also influence the coordinates of atoms in other bonds that link to the rotated node of the bond, resulting in a change to the gesture of the whole ligand. Reducing complexity by replacing the torsion operation with the perturbation operation in Euclidean space makes the model easier to train and more efficient in inference.

### 2.4 Model Architecture

The score model **s**(**x**_*l*_, **x**_*p*_, **h**_*l*_, **h**_*p*_, *t*) is constructed to take as input the 3D coordinates **x**_(·)_ and type **h**_(·)_ for all the heavy atoms of ligand *l* and protein pocket *p*, along with diffusion time *t*. The output of the score model predicts the translation score and rotation score in the tangent space *T*_**r**_ 𝕋 (3)⊕*T*_***ω***_ *SO*(3) and perturbation score in Euclidean space R^3*n*^. The tangent space of translation and rotation is isomorphic to ℝ^6^, where *T*_**r**_ 𝕋 (3) ≅ ℝ^3^ corresponding to the translation vector and *T*_***ω***_ *SO*(3) ≅ R^3^ corresponding to the rotation Euler vector. Model components are described below and detailed in the Model Architecture section.

As shown in Fig. 1(a), a ligand encoder E_*l*_ and a protein pocket encoder ℰ _*p*_ with the same SE(3)-invariant transformer architecture (Zhou et al., 2022) are used to encode the input ligand and protein representations **y**_*s*_ = ℰ _*s*_(**x**_*s*_, **h**_*s*_), *s* = *l* or *p*. Since the coordinates of the molecule are not *SE*(3)-invariant quantities, a distance matrix is used to encode the position of atoms, which is *SE*(3)-invariant (see detail in the Appendix Proof section). In addition, an edge-type matrix defined by the type of atom pair of each edge is used to encode the potential bond information. A 3D spatial positional encoding matrix is then generated from the distance and edge type matrix by a pair-type aware Gaussian kernel (Shuaibi et al., 2021), following Zhou et al. (2022). The ligand and protein representation are then concatenated **y** = **y**_*l*_ | **y**_*p*_ and fed into the decoder network 𝒟, along with the cross pair representation between ligand and protein pocket to get the score (See detail in the Decoder Network section). For the decoder network 𝒟, as shown in Fig. 1(b), we designed a scalable new SE(3)-equivariant transformer architecture (green part in Fig. 1(b)), differing from previous work (Chatzipantazis et al., 2022). Inspired by the gate control mechanism in LSTM (Graves and Graves, 2012), we separate the SE(3)- invariant components and SE(3)-equivariant components into two branches, while allowing interaction between these two branches through dot-product (See detail in the Decoder Network section). The concatenate representation from the final decoder layer was then used to produce the system diffusion score (translation and rotation) and the atom-wise diffusion score (perturbation score). To produce the system diffusion score in ℝ^6^, we take the first representation of output layer, which corresponds to the padding token *< s >*. Producing atom-wise score is quite straightforward with the use an atom-wise projection.

**Fig. 1.**
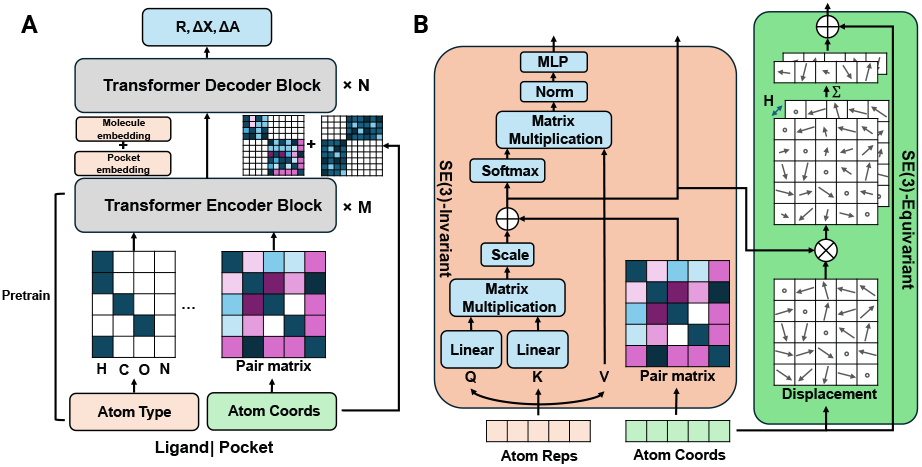
The architecture of the score network in DTMol. (A). The score network consists of two pre-trained encoder networks and a decoder network. The encoder networks individually processes the ligand and the protein pocket, generating encoded embeddings that are subsequently used by a decoder network to predict scores. Differ from the usual transformer, the encoder network features a *SE*(3)-invariant attention layer that maintains a pair representation, which is initially created by a Gaussian kernel on the distance matrix and the edge-type tensor. (B) The decoder block includes the same *SE*(3)-invariant branch as in the encoder block (orange part), and a unique *SE*(3)-equivariant branch that updates the coordinates by modifying the displacement tensor of the input coordinates through tensor-producting with the pair representation (green part).

### 2.5 Model Training

Our model training includes two stages: a sampling stage and a score matching stage. In the *sampling* stage, perturbation and rototranslation noise are sampled and applied to the ligand, while only perturbation noise is applied to the protein pocket. The model is then trained to minimize the score loss given the noised ligand and protein pocket.

#### Algorithm 1

Model Training Procedure

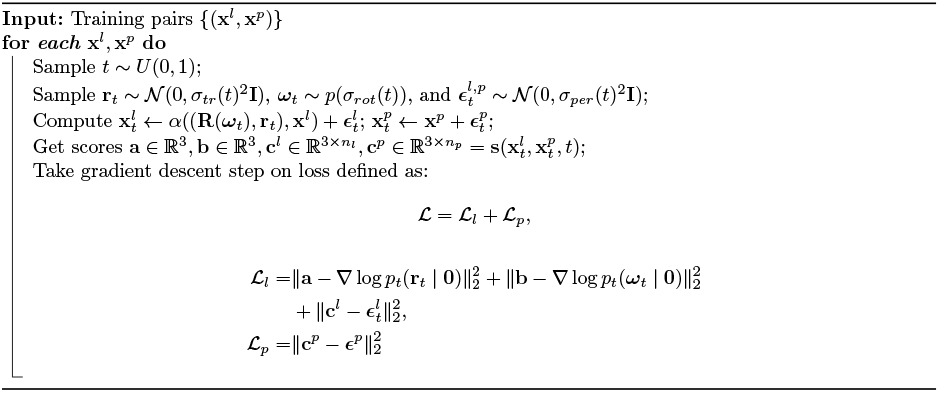

Several noise schedulers *σ*(*t*) have been tested (Nichol and Dhariwal, 2021; Corso et al., 2023; Leach et al., 2022), and we found no significant difference among the noise schedulers used, except for the linear one, which increases the noise too quickly and makes the model hard to train. We adopted a warmup strategy (Loshchilov and Hutter, 2016) that begins with a small learning rate. We found it vital for the model to converge. Additionally, we freeze the pretrained encoder models until a certain number of epochs has been reached (see the Training Details section for more information).

## 3 Experiments

We evaluate our models on the PDBBind (Liu et al., 2017) and PoseBuster datasets (Buttenschoen et al., 2024), comparing to other machine learning models and traditional search-based models. We demonstrated the ability of our models on a virtual scrrening task on Drugbank molecule dataset, with wet-lab assay validation.

### 3.1 Experimental Setup

We preprocessed the PDBBind v2020 dataset (Liu et al., 2017) by removing the duplicate entries and selecting the protein-ligand pairs with resolution less than 2.5 Å and R-free less than 0.25. As DTMol requires the protein pocket as input, we extracted the residues around the true binding position (see detail in the Dataset section). For a fair comparison, the same information was exposed to other models, by sitting the initial ligand position at the center of the protein pocket. The performance of molecular docking methods was evaluated by using the Top-5 and Top-1 RMSD, where multiple docking simulations were conducted for each protein and the best one or five results reported by the model were selected for comparison to the reference ligand pose. We applied the smina as our confidence model to rank our generated conformations. Polar hydrogen atoms are inferred and added back to the structure using RDKit in order to obtain binding affinity scores with smina. The proportion of generated conformations with RMSD below a certain threshold was calculated.

### 3.2 Performance on docking dataset

Our results demonstrate that our model outperforms both search-based and machine learning-based docking methods across all evaluated datasets (Tables 1 and 2). Notably, models generally exhibit much better performance on the PoseBuster benchmark dataset (Table 2) and PoseBuster Astex Diverse set (Table 3) compared to the PDBBind dataset (Table 1). This difference is likely due to the time-split strategy used in PDBBind dataset to get the test split, which aims to prevent information leakage from the test set to the training set. The superior performance on the two PoseBuster datasets may indicate potential information leak, highlighting the importance of rigorous data splitting techniques in evaluating model performance.

**Table 1.** Docking Results for PDBBind test set (363 complexes)

| Methods | Top-1 RMSD |  |  |  | Top-5 RMSD |  |  |  |
| --- | --- | --- | --- | --- | --- | --- | --- | --- |
|  | %<1Å | %<2Å | %<5Å | Med. | %<1Å | %<2Å | %<5Å | Med. |
| Diffdock* | 12.50 | 28.93 | 57.86 | 4.37 | **26.07** | 47.86 | 74.64 | 2.158 |
| Smina | 12.17 | 26.64 | 51.97 | 4.690 | 15.79 | 43.42 | 72.70 | 2.450 |
| Gnina | 12.83 | 36.51 | 62.17 | 3.349 | 16.78 | 50.33 | 77.63 | 1.982 |
| DiffBindFR | 8.55 | 33.22 | 70.65 | 3.179 | 14.80 | 48.36 | **85.20** | 2.070 |
| DTMol | **16.72** | **45.05** | **74.54** | **3.174** | 20.28 | **52.36** | 79.63 | **1.879** |

**Table 2.** Docking Results for PoseBuster (428 complexes)

| Methods | Top-1 RMSD |  |  |  | Top-5 RMSD |  |  |  |
| --- | --- | --- | --- | --- | --- | --- | --- | --- |
|  | %<1Å | %<2Å | %<5Å | Med. | %<1Å | %<2Å | %<5Å | Med. |
| Diffdock* | 11.30 | 22.61 | 48.12 | 5.550 | 24.93 | 42.32 | 69.57 | 2.599 |
| Smina | 32.55 | 50.47 | 76.42 | 1.961 | 41.98 | 71.22 | 92.69 | 1.182 |
| Gnina | 38.55 | 65.65 | 90.42 | 1.372 | 48.83 | **81.07** | 96.96 | 1.037 |
| DiffBindFR | 14.02 | 38.55 | 78.27 | 2.627 | 23.60 | 57.01 | 93.46 | 1.745 |
| DTMol | **43.16** | **77.65** | **94.04** | **1.253** | **49.32** | 79.04 | **97.53** | **1.025** |

**Table 3.**
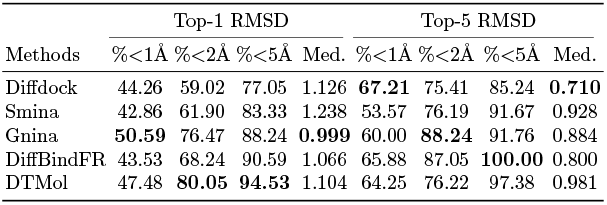
Docking Results for PoseBuster Astex (85 complexes)

| Methods | Top-1 RMSD |  |  |  | Top-5 RMSD |  |  |  |
| --- | --- | --- | --- | --- | --- | --- | --- | --- |
|  | %<1Å | %<2Å | %<5Å | Med. | %<1Å | %<2Å | %<5Å | Med. |
| Diffdock | 44.26 | 59.02 | 77.05 | 1.126 | **67.21** | 75.41 | 85.24 | **0.710** |
| Smina | 42.86 | 61.90 | 83.33 | 1.238 | 53.57 | 76.19 | 91.67 | 0.928 |
| Gnina | **50.59** | 76.47 | 88.24 | **0.999** | 60.00 | **88.24** | 91.76 | 0.884 |
| DiffBindFR | 43.53 | 68.24 | 90.59 | 1.066 | 65.88 | 87.05 | **100.00** | 0.800 |
| DTMol | 47.48 | **80.05** | **94.53** | 1.104 | 64.25 | 76.22 | 97.38 | 0.981 |

### 3.3 Validation on a virtual screening task

We conducted a virtual screening task using DTMol, in which 9,137 drugs were screened from the DrugBank database v5.1.8 to find potential candidates that bind to the Janus Kinase (JAK2) protein. A complex structure with PDB ID 6BBV was used as the target protein, and the location of the binding pocket was defined by the ligand D7D in the complex. The ligand binding pose was predicted by DTMol (Appedix Fig. S1), and a score was then given by smina (see the Appendix Evaluation section for additional details). Candidate molecules were ranked by the score, and the top 20 molecules were selected for further experimentation. We also applied the physics-based docking software Autodock Vina to screen the same database and included the top 20 molecules as well. A total of 27 out of 40 molecules were successfully tested using Time-Resolved Fluorescence Resonance Energy Transfer (TR-FRET) competitive binding assay to determine the activity score under a 100 nM concentration. Two known JAK2 inhibitors Ruxolitinib and Tofacitinib are added into the assay experiment as the control sample under the same condition. Then 15 molecules with the highest activity scores from the first round were selected for a second round of activity assays at 50 nM and 100 nM concentrations (Fig. 2). Molecules that demonstrated higher activity scores than both known JAK2 inhibitors at both concentration levels were considered potential JAK2 inhibitors. All tests were conducted in a blind setting, where each molecule was identified only by a numerical index, with the identities revealed only after the whole experiments were completed. DTMol achieved a higher Spearman’s rank-correlation score (0.192) than Autodock (−0.146) between the experimental ligand inhibition rank and predicted rank. Additionally, two new compounds from the AI condidate list were identified as potential JAK2 inhibitor (Table S2).

**Fig. 2.**
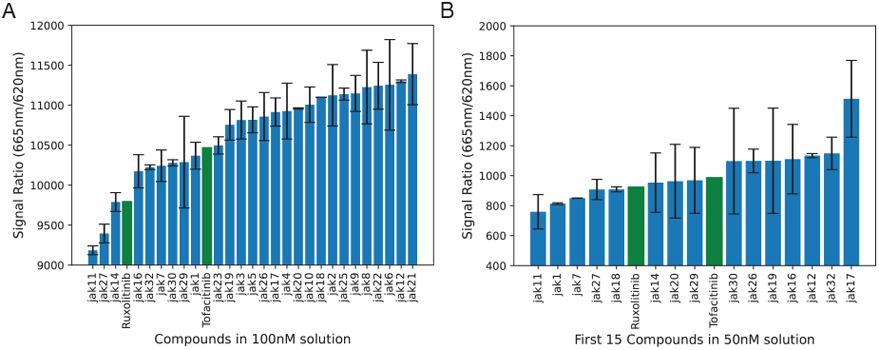
TR-FRET assay results for 27 compounds at 100nM (A) and 15 compounds at 50nM (B). A lower signal ratio indicates stronger inhibition by the compound. Green bars indicate the two known JAK2 inhibitors, Ruxolitinib and Tofacitinib, which are used as controls.

## 4 Experimental Details

### 4.1 Dataset

The PDBBind dataset is a comprehensive collection of protein-ligand complexes with known 3D structures and binding affinities, widely used for developing and evaluating computational methods for predicting protein-ligand interactions. The latest version, PDBBind v2020, contains 19,120 complexes. We use a time split strategy same as Stärk et al. (2022), which divides the dataset into training and test sets based on the deposition date of the complexes in the Protein Data Bank (PDB), where 363 complexes deposit later than 2019 are selected as test dataset, mimicking a real-world scenario where a model is trained on historical data and applied to new, unseen complexes. The PoseButser dataset is another collection of 513 diverse protein-ligand complexes designed for evaluating the accuracy of docking poses, including 85 diverse protein-ligand set (PoseBuster Astex) and 428 recent protein-ligand complexes released from 2021 onwards. Because the PDBBind training dataset includes only protein–ligand complexes deposited before 2019, there is no overlap with the PoseBuster test set. On the other hand, the PoseBuster Astex dataset contains structures from the PDB-bind training dataset, so it is not suitable to serve as a benchmark dataset. However, as it contains drug-like molecules of pharmaceutical or agrochemical interest, we still report the score on it. Although a time-split does not guarantee complete removal of sequence or structural homologs, both PDBBind and PoseBuster employ strategies to reduce redundancy (e.g., avoiding near-identical protein by restricting sequence similarity to ¡90%). Furthermore, we checked for any obvious overlap between our training data and the PoseBuster test set and found no exact duplicates or near-identical entries.

### 4.2 Vitrual Screening Task

We selected the Janus Kinase 2 (JAK2) protein as our target for virtual screening due to its crucial role in the JAK-STAT signaling pathway and its involvement in various human diseases, including several types of cancer such as breast cancer, prostate cancer, and leukemia. Additionally, JAK2 has several recently identified inhibitors that are not in our training dataset collection. We selected the top 20 ranked drugs from each of the docking results obtained using DTMol plus the smina scoring function and Autodock Vina, and purchased all available drugs. Fifteen drugs out of the 40 top-ranked candidates could not be acquired due to supplier shortages or restrictions. We also added two known inhibitors as control samples to validate the efficiency of our experimental protocol. Finally, 27 drugs (25 top-ranked candidates and 2 control samples) were successfully tested, and a rank was acquired from the activity assay experiment. This rank was then compared with the rank given by the docking model.

### 4.3 Training Details

We adopted a learning rate scheduler with a warmup strategy which sets up a small initial learning rate (1e-6) and then increases gradually. Several schedulers (Loshchilov and Hutter, 2016) have been tested and a Linear scheduler has been chosen in our case. The pretrained encoder was frozen in the first 200 epochs and then finetuned along with the decoder for the rest of the training process. The model was trained on two RTX A100 80GP GPUs for 500 epochs (around 7 days). For inference, the model can be run on a single RTX 4090 24 GB GPU. For every 100 training steps, we run a validation step on a random batch from a separate validation set to get the perturbation, rotation, and translation score loss. For every 5 epochs, we run inference of the model with 20 reverse diffusion steps through a separate test dataset with 300 complexes to get the RMSD score.

#### Hyperparameters

We have tested several hyperparameters and found that some are important to model convergence and performance, while others have little effect. All the relevant hyperparameters are highlighted in Table S1. Meanwhile, we found that DTMol is insensitive to several previously reported parameters, including the number of sampling steps during inference. We did not find a consistent performance difference when changing the sampling steps from 20 to 40 across datasets. However, we found that a much larger number of sampling steps, such as 500, hurt the model performance on all datasets, consistent with a previous study by Corso et al. (2023). We also found that increasing the number of warmup rounds from 5 had little effect on model convergence and did not result in a lower training loss, so we use 5 warmup rounds. However, we did find that different learning rate schedulers after the warmup rounds are important for model convergence. We discovered that the Cosine-style Scheduler prevented the model from converging and even broke a converged model, increasing the training loss of the model, similar to catastrophic forgetting. A linear learning rate scheduler, on the other hand, did well and is quite robust. Other important hyperparameters include the maximum variance added to the translation score, as a small value would restrict the model’s ability to recover ligands from longer distance, but a large value would increase the variation of the sampled input thus making the training process more variant and harder. We have not tested a switching strategy that dynamically changes this maximum variance of noise added, which could be an interesting future work.

## 5 Conclusion

We present DTMol, a diffusion transformer model with a novel *SE*(3) equivarinat attention layer for pocket-based molecular docking. Attention-based transformer neural network uses attention mechanism to transfer information between nodes instead of the neighbourhood graph in the graph neural network, allowing more efficient information flow. However, computing the attention matrix is equivalent to performing a forward pass on a complete graph in a graph neural network, incurring a quadratic cost in the number of atoms and thus requiring substantial memory for a full-atom model. Other transformer variants with a memory-efficient attention mechanism can be further explored to achieve a full protein docking model. A full protein docking model would help address allostery-driven ligand binding, in which the protein pocket dynamically adapts its conformation in response to allosteric signals. We believe training on all heavy atoms is necessary and benefits the model’s performance on docking tasks. Diffusion models trained on only alpha-carbon atoms, such as Diffdock, demonstrate difficulty in matching conventional docking methods for pocket-based docking tasks (Jain et al., 2024). It would be an interesting future direction to see if, without a pretrained encoder, the decoder-only model is sufficient. Another limitation is the availability of the dataset, as it is hard to obtain exact crystal structures of protein-ligand complexes. Use of synthesized data or other data augmentation techniques can be tested to improve the model’s generalization ability. Adding more diverse holo pocket structures to the training dataset would also reduce the bias to cognate ligands for the pocket when conducting docking using the trained model, which could be a future direction.

## 6 Code Availability

The source code for DTMol is available at https://github.com/haotianteng/dtmol. Additionally, the benchmark code can be accessed at https://github.com/haotianteng/MolecularDocking.

## 7 Acknowledgements

This work was supported in part by the US National Science Foundation [III-2232121], the US National Institutes of Health [R01HG012470], the SJTU “Star of Jiaotong University” program, Medical Engineering Cross Research Funding [YG2023QNA13]. H.T., R.W., Y.Y., and C.K. conceived the study.

H.T. designed the DTMol model, and H.T. and Y.S. derived the diffusion form. H.T. implemented and tested multiple variants of the DTMol model. H.T. conducted the benchmarking. R.W. performed the JAK2 assay experiment. H.T. and Y.S. wrote the initial draft, and H.T., Y.S., and C.K. refined the manuscript. We used ChatGPT to correct grammatical errors and improve the flow of early drafts of this manuscript. We thank Euxhen Hasanaj for the helpful discussion in the initial of this project. We thank David Koes from University of Pittsburgh for the helpful discussion on the virtual screening task and evaluation.

## Appendix

### 1 Details of Model Architecture

The positions and types of all the heavy atoms in the ligand and protein pocket are encoded by a ligand encoder E_*l*_ and a protein pocket encoder E_*p*_, respectively. The ligand and protein pocket embeddings **y**_*l*_ and **y**_*p*_ are then concatenated and fed into the decoder (score network), along with the ligand and protein coordiantes **x**_*l*_ and **x**_*p*_, to predict the score of the diffusion process at time *t, s*(**y**_*l*_, **y**_*p*_, **x**_*l*_, **x**_*p*_, *t*). The decoder consists of parallel SE(3)-invariant and SE(3)-equivariant transformer layers, where the final outputs from the two branches are dot-producted to generate the final score prediction. The detailed architecture of the encoders and the score network is as follows.

#### 1.1 Transformer encoder with pair representation attention

A pair representation **P** is added to each attention matrix in the transformer encoder and is updated in each attention layer. An initial pair representation for a molecule (ligand or pocket) is generated by a distance matrix **D** ∈ ℝ^*n*×*n*^ and a one hot edge type matrix **E** ∈ {0, 1}^*n*×*n*×*m*^, where *n* is the number of atoms and *m* is the number of edge types. The distance matrix is calculated by the Euclidean distance between each pair of atoms in the molecule. Each edge type is defined by the pair of atoms it connects, resulting in a total of *m* = *c*^2^ possible edge types, where *c* is the number of distinct atom types. For instance, if a carbon atom and an oxygen atom are indexed as 3 and 4, respectively, in the atom dictionary, then an edge connecting these two atoms results in a one-hot vector with a one in the entry 3*c* + 4.

The distance matrix and the edge type matrix are then used to create the initial pair representation **P** ∈ ℝ^*n*×*n*×*h′*^, by the Gaussian kernel 𝒢 (Shuaibi et al., 2021):

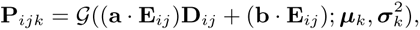

where **a** ∈ R^*m*^ and **b** ∈ R^*m*^ are learnable embedding vectors, ***µ*** ∈ ℝ^*h′*^ and ***σ*** ∈ ℝ^*h′*^ are learnable mean and variance of the Gaussian kernel, where *h*^′^ is the number of kernel basis. A linear projection **U** ∈ ℝ^*h′*×*h*^ is then multiplied with **P**, generating **P**^1^ ∈ ℝ^*n*×*n*×*h*^ which is the input of the first layer of the transformer architecture. The attention matrix for each layer is then computed as:

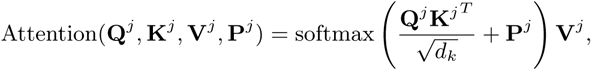

where *j* represents the *j*-th layer, and **Q**^*j*^, **K**^*j*^, **V**^*j*^ are trainable matrices. The pair representation **P**^*j*^ will then be updated as:

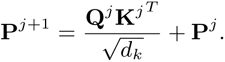

#### 1.2 Decoder Network

The decoder network D takes the concatenated ligand and protein representations **y** = **y**_*l*_ | **y**_*p*_, along with the cross pair representation between ligand and protein pocket, to predict the score of the diffusion process. The cross pair representation 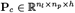 is generated using the same Gaussian kernel method metioned above, with cross distance matrix and cross edge type matrix. A full pair representation matrix 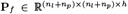 is then constructed by combining the ligand pair representation matrix **P**_*l*_ and the protein pocket pair representation matrix **P**_*p*_ with the cross pair representation matrix **P**_*c*_ by:

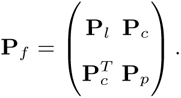

In addition to the full pair representation matrix **P**_*f*_ and the concatenated ligand and protein representations **y**, decoder network also takes the coordinates of ligand **x** as input. A normalized displacement matrix 𝔻 ∈ ℝ^*n*×*n*×3^ is calculated as:

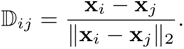

To create the SE(3)-equivariant displacement tensor **S** ∈ ℝ^*n*×*n*×*h*×3^, we apply the tensor product — implemented using the *e*3*nn* library (Geiger and Smidt, 2022)— to the displacement matrix D along with the initial pair representation **P**^1^

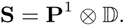

**S** is then used to update the coordinates **x** at each head:

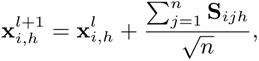

while the initial **x**_*h*_ for each head is just a copy of the input coordinates **x**. The process is repeated at each layer, where a new normalized displacement matrix is calculated given the updated coordinates, and new SE(3)-equivariant displacement tensor is generated by dot-producting the displacement matrix 𝔻^*l*^ ∈ ℝ^*n*×*n*×*h*×3^ and pair-representation **P**^*l*^ (tensor product is only used in the first layer where displacement matrix has no head dimension).

#### 1.3 Output layers

Final score prediction is given by a dot-product of the final representation of the SE(3)-invariant branch and the SE(3)-equivariant branch, given by:

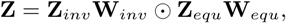

where **W**_*inv*_ ∈ ℝ^*hd*×*o*^ and **W**_*equ*_ ∈ ℝ^3*h*×*o*^ are the projection matrices that project the input to desired output dimension o, **Z**_*inv*_ ∈ ℝ^*n*×*h*×*d*^ is the output of final attention layer following a MLP, and **Z**_*equ*_ ∈ R^*n*×*h*×3^ is given by:

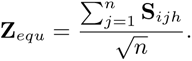

To predict the translation and rotation score, a two-layer MLP with GeLU activation function is used to get **Z**_*inv*_, taking the first representation in the sequence of the attention output, which corresponds to a padded special token ¡s¿, and projecting it to a length-6 vector. As for the atom-wise diffusion score, another two-layer MLP with GeLU activation function is used to get **Z**_*inv*_, taking the full sequence of the attention layer output, resulting the perturbation score prediction 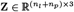.

### 2 Proofs

#### 2.1 Proof for distance matrix being *SE*(3)-invariant

##### Lemma 1.

*Let* **x** ∈ ℝ^3*n*^ *be the position coordinates of all n atoms in a molecule, and let* **D** ∈ ℝ^*n*×*n*^ *be the distance matrix such that* **D**_*ij*_ = ||**x**_*i*_ − **x**_*j*_ ||_2_. *For any element g* ∈ *SE*(3), *representing a rototranslational operation in 3D Euclidean space, if we apply g to x to obtain new coordinates x*^′^ *such that* 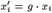 *for any atom i, and a new distance matrix* **D**^′^ *such that* 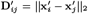, *then we have* **D**^′^ = **D**.

*Proof*. Since each roto-translational operation *g* on a 3D coordinate *x*_*i*_ can be represented by an affine transformation *Rx*_*i*_ + *v*, where *R* ∈ *SO*(3) is a rotational matrix and *v* ∈ ℝ^3^ is a translational vector, we have:

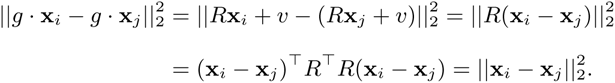

The last equality holds since for each element *R* in *SO*(3), we have *R*^⊤^*R* = *RR*^⊤^ = **I**. Therefore, for each *i, j* pair, we have:

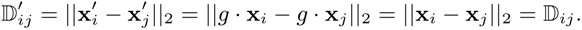

#### 2.2 Proof for coordinate update strategy being *SE*(3)-equivariant

##### Lemma 2.

*Let* 𝔻 ∈ ℝ^*n*×*n*×3^ *be the normalized displacement tensor such that* 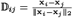. *For any element g* ∈ *SE*(3), *if we apply g to x to obtain new coordinates x*^′^, *for any transformation Ψ* (**x**) = **x** + _*j*_ *Φ*(**x**)_*ij*_ 𝔻_*ij*_ *with Φ*((*x*)) *being SE*(3)*-invariant transformation, the following hold Ψ* (*α*_*g*_ (**x**)) = *α*_*g*_ (*Ψ* (**x**)), *i*.*e. Ψ* (**x**) *is a SE*(3)*-equivariant transformation*.

*Proof*. As *α*_*g*_ (**x**_*i*_) = *R***x**_*i*_ + *v*, we have:

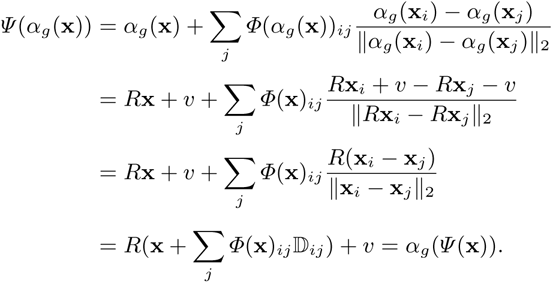

#### 2.3 Proof of group action under expectation

Let **x** = {**x**_1_, **x**_2_, …, **x**_*n*_} be the position coordinates of all *n* atoms. We can apply the following operations during the diffusion process:

- System Rotation around the center 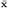

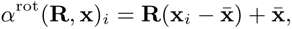

where **R** ∈ *SO*(3) is a rotation matrix.
- System Translation

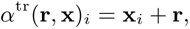

where **r** ∈ ℝ^3^ is a translation vector.
- Perturbation Noise

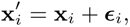

where ***ϵ***_*i*_ ∼ 𝒩 (**0**, *σ*^2^**I**) are independent zero-mean Gaussian noises.

To prove the operations forms a group action, we need to prove that the combined operation *α*, defined as:

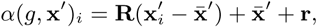

with *g* = (**R, r**), forms a group action on the noised point cloud **x**^′^.

*Proof*. First, note that the noise has zero mean:

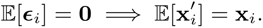

The expected center of the perturbed point cloud keeps the same as original point cloud:

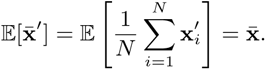

To prove *α* forms a group action under expectation, we need to prove the identity and the compatibility property of */alpha*.

**Proof of identity**: Let *e* = (**I, 0**) be the identity element. Then:

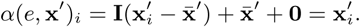

Taking expectation:

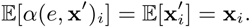

Thus, the identity property holds in expectation.

**Proof for Compatibility Property**:

Let *g*_1_ = (**R**_1_, **r**_1_) and *g*_2_ = (**R**_2_, **r**_2_). Their composition is:

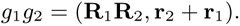

We need to show:

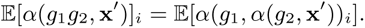

Compute the left-hand side (LHS):

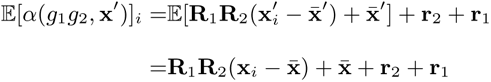

Compute the right-hand side (RHS):

First, compute 𝔼 [*α*(*g*_2_, **x**^′^)]_*i*_:

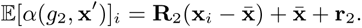

The expected center after applying *g*_2_ is:

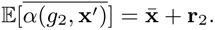

Then:

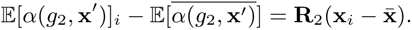

Now, compute the RHS:

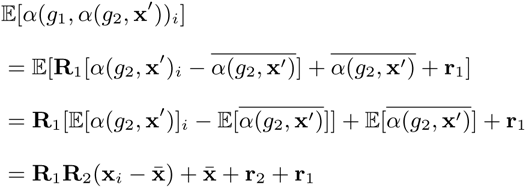

Comparing LHS and RHS:

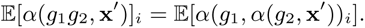

Thus, the compatibility property holds in expectation.

#### 2.4 Diffusion form of SMLD and DDPM

Below we provide a straight-forward derivation for SMLD and DDPM diffusion process. For forward diffusion process in SMLD with Gaussian kernel *P* (**x**_*i*_|**x**_*i*−1_) = 𝒩 (**x**_*i*_; **x**_*i*−1_, (*σ*^2^ − *σ*^2^)**I**)

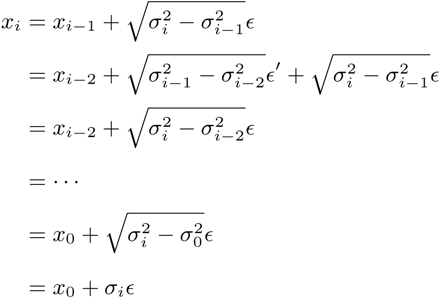

If we assume *σ*_0_ = 0, and notice for two independent gaussian variable 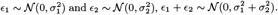.

Similarly, for forward diffusion process in DDPM with Gaussian kernel 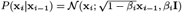

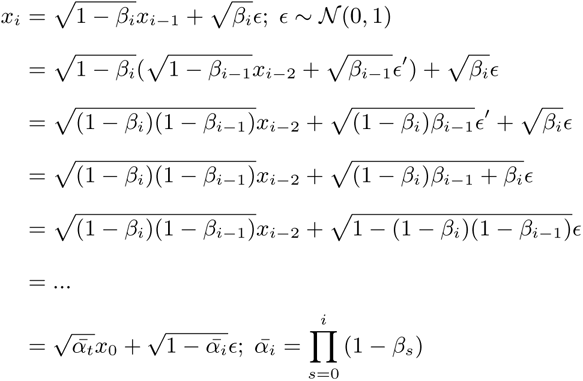

### 3 Evaluation Details

We compared DTMol with two search-based docking methods Smina and gnina, and with two previously reported high-accurate machine-learning based docking methods Unimol and DiffDock.

*SMINA* Koes et al. (2013) uses Autodock-Vina docking kernel with a new scoring function. We set “exhaustiveness” to 100 (default is 8) for better performance, and set “num modes=10” to generate 10 candidates. Other settings are kept as default.

*GNINA* McNutt et al. (2021) equipped with an additional 3D CNN for scoring function. Same to Smina, we set the ‘–exhaustiveness 100’ and ‘–num modes’ to 10.

*Unimol* Zhou et al. (2022) is a universal molecular embedding model for multiple downstream tasks, including molecular docking. We directly use the weights pretrained on binding pose prediction provided by Unimol, and all parameters are kept as default. For a fair comparison, all pockets are the same according to different pocket-based docking models.

*DiffDock* Corso et al. (2023) is a generative diffusion model designed for blind docking. As it is not able to specify a protein pocket with DiffDock, we generated 50 samples on the whole protein with DiffDock, and then selected the 10 samples with minimum RMSD as the final collection. We calculate top1 and top5 scores among them according to the score given by the confidence model.

*DiffBindFR* Zhu et al. (2024) is another generative diffusion model designed for pocket-based molecular docking. Similar to DiffDock, it employs a graph neural network with a torsion diffusion process. Additionally, the model considers protein side chain flexibility and allows specifying a protein pocket with a reference ligand as the starting position for the reverse diffusion process (docking). We used the default settings of the model, generating 40 samples for each protein-ligand pair, and used the Smina scoring function to rank the predictions.

*DTMol* run with Smina as a scoring function. Multiple samples (default is 10) are generated by DTMol for each docking process, which are then scored and ranked by Smina.

**Table S1.**
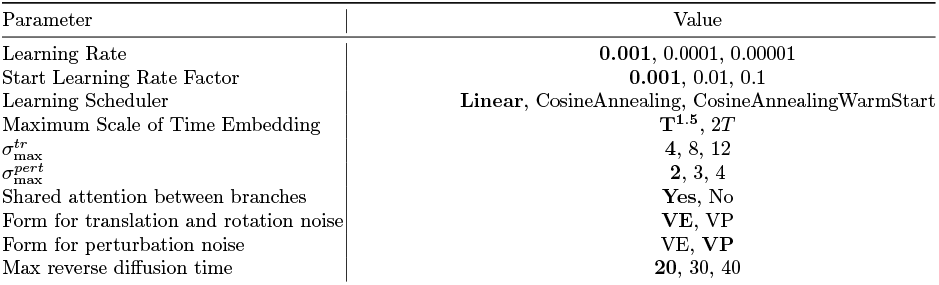
Hyperparameters searched for the DTMol model; the final parameters are highlighted in bold.

**Supplementary Figure S1.**
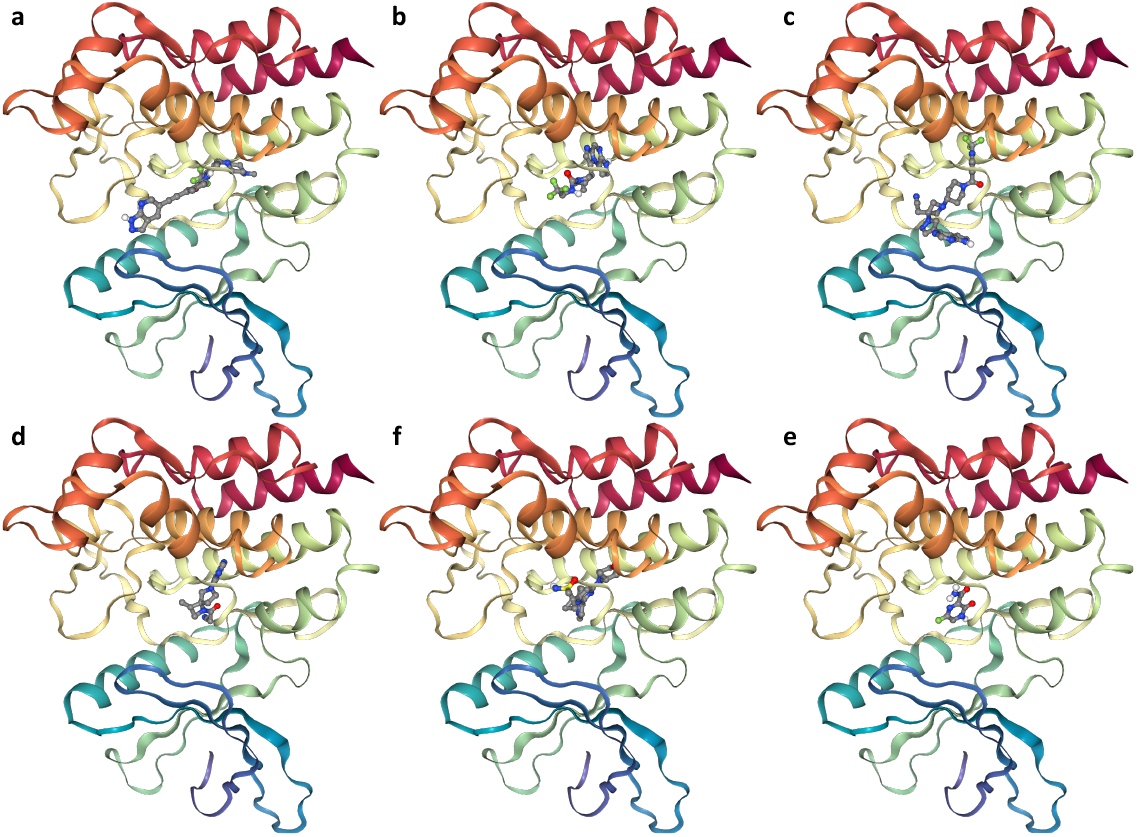
An illustration of docked poses generated by DTMol for six drugs binding to the JAK2 protein (PDB ID: 6bbv): (a) HQP1351, (b) Upadacitinib, (c) Itacitinib, (d) Delgocitinib, (e) Ceralascrtib, and (f) Favipiravi.

**Table S2.**
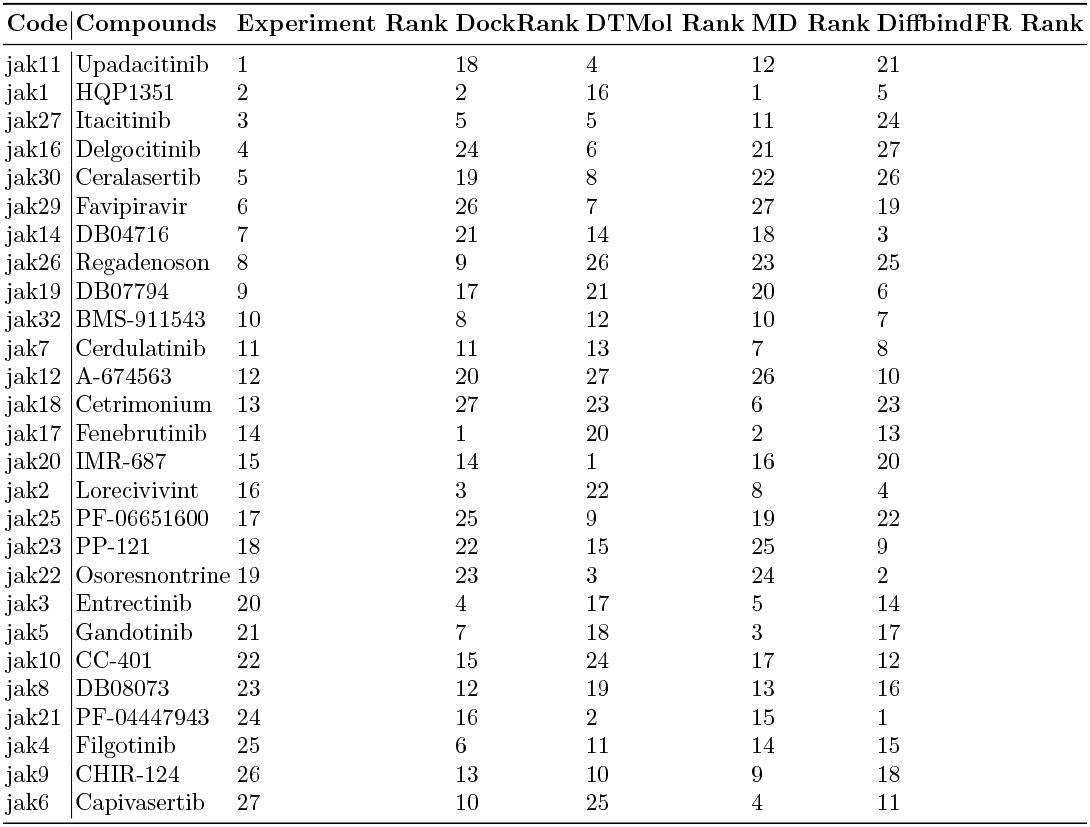
Comparison of the ranks given by the assay experiment (Experiment Rank), docking (DockRank), DTMol (AI Rank), and docking followed by MD relaxation (MD Rank) for different compounds. DTMol got the only positive Spearman’s rank correlation (0.192), while the DiffbindFR, the conventional docking method and conventional docking followed by molecular dynamic relaxation got negative correlation scores (−0.264, −0.146 and −0.194)

