## Appendix for "DTMol: Pocket-based Molecular Docking using Diffusion Transformers"

### 1 Details of Model Architecture

The positions and types of all the heavy atoms in the ligand and protein pocket are encoded by a ligand encoder $\mathcal{E}_l$ and a protein pocket encoder $\mathcal{E}_p$, respectively. The ligand and protein pocket embeddings $\mathbf{y}_l$ and $\mathbf{y}_p$ are then concatenated and fed into the decoder (score network), along with the ligand and protein coordiantes $\mathbf{x}_l$ and $\mathbf{x}_p$, to predict the score of the diffusion process at time $t$, $s(\mathbf{y}_l, \mathbf{y}_p, \mathbf{x}_l, \mathbf{x}_p, t)$. The decoder consists of parallel SE(3)-invariant and SE(3)-equivariant transformer layers, where the final outputs from the two branches are dot-producted to generate the final score prediction. The detailed architecture of the encoders and the score network is as follows.

#### 1.1 Transformer encoder with pair representation attention

A pair representation $\mathbf{P}$ is added to each attention matrix in the transformer encoder and is updated in each attention layer. An initial pair representation for a molecule (ligand or pocket) is generated by a distance matrix $\mathbf{D} \in \mathbb{R}^{n\times n}$ and a one hot edge type matrix $\mathbf{E} \in \{0,1\}^{n\times n\times m}$, where $n$ is the number of atoms and $m$ is the number of edge types. The distance matrix is calculated by the Euclidean distance between each pair of atoms in the molecule. Each edge type is defined by the pair of atoms it connects, resulting in a total of $m = c^2$ possible edge types, where $c$ is the number of distinct atom types. For instance, if a carbon atom and an oxygen atom are indexed as 3 and 4, respectively, in the atom dictionary, then an edge connecting these two atoms results in a one-hot vector with a one in the entry $3c + 4$.

The distance matrix and the edge type matrix are then used to create the initial pair representation $\mathbf{P} \in \mathbb{R}^{n\times n\times h'}$, by the Gaussian kernel $\mathcal{G}$ (Shuaibi et al., 2021):

$$\mathbf{P}_{ijk} = \mathcal{G}((\mathbf{a}\cdot\mathbf{E}_{ij})\mathbf{D}_{ij} + (\mathbf{b}\cdot\mathbf{E}_{ij}); \boldsymbol{\mu}_k, \boldsymbol{\sigma}_k^2),$$

where $\mathbf{a} \in \mathbb{R}^m$ and $\mathbf{b} \in \mathbb{R}^m$ are learnable embedding vectors, $\boldsymbol{\mu} \in \mathbb{R}^{h'}$ and $\boldsymbol{\sigma} \in \mathbb{R}^{h'}$ are learnable mean and variance of the Gaussian kernel, where $h'$ is the number of kernel basis. A linear projection $\mathbf{U} \in \mathbb{R}^{h'\times h}$ is then multiplied with $\mathbf{P}$, generating $\mathbf{P}^1 \in \mathbb{R}^{n\times n\times h}$ which is the input of the first layer of the transformer architecture. The attention matrix for each layer is then computed as:

$$\text{Attention}(\mathbf{Q}^j, \mathbf{K}^j, \mathbf{V}^j, \mathbf{P}^j) = \text{softmax}\left(\frac{\mathbf{Q}^j{\mathbf{K}^j}^T}{\sqrt{d_k}} + \mathbf{P}^j\right)\mathbf{V}^j,$$

where $j$ represents the $j$-th layer, and $\mathbf{Q}^j, \mathbf{K}^j, \mathbf{V}^j$ are trainable matrices. The pair representation $\mathbf{P}^j$ will then be updated as:

$$\mathbf{P}^{j+1} = \frac{\mathbf{Q}^j {\mathbf{K}^j}^T}{\sqrt{d_k}} + \mathbf{P}^j.$$

#### 1.2 Decoder Network

The decoder network $\mathcal{D}$ takes the concatenated ligand and protein representations $\mathbf{y} = \mathbf{y}_l \mid \mathbf{y}_p$, along with the cross pair representation between ligand and protein pocket, to predict the score of the diffusion process. The cross pair representation $\mathbf{P}_c \in \mathbb{R}^{n_l \times n_p \times h}$ is generated using the same Gaussian kernel method metioned above, with cross distance matrix and cross edge type matrix. A full pair representation matrix $\mathbf{P}_f \in \mathbb{R}^{(n_l+n_p)\times(n_l+n_p)\times h}$ is then constructed by combining the ligand pair representation matrix $\mathbf{P}_l$ and the protein pocket pair representation matrix $\mathbf{P}_p$ with the cross pair representation matrix $\mathbf{P}_c$ by:

$$\mathbf{P}_f = \begin{pmatrix} \mathbf{P}_l & \mathbf{P}_c \\ \mathbf{P}_c^T & \mathbf{P}_p \end{pmatrix}.$$

In addition to the full pair representation matrix $\mathbf{P}_f$ and the concatenated ligand and protein representations $\mathbf{y}$, decoder network also takes the coordinates of ligand $\mathbf{x}$ as input. A normalized displacement matrix $\mathbb{D} \in \mathbb{R}^{n \times n \times 3}$ is calculated as:

$$\mathbb{D}_{ij} = \frac{\mathbf{x}_i - \mathbf{x}_j}{\|\mathbf{x}_i - \mathbf{x}_j\|_2}.$$

To create the SE(3)-equivariant displacement tensor $\mathbf{S} \in \mathbb{R}^{n \times n \times h \times 3}$, we apply the tensor product — implemented using the *e3nn* library (Geiger and Smidt, 2022)— to the displacement matrix $\mathbb{D}$ along with the initial pair representation $\mathbf{P}^1$

$$\mathbf{S} = \mathbf{P}^1 \otimes \mathbb{D}.$$

$\mathbf{S}$ is then used to update the coordinates $\mathbf{x}$ at each head:

$$\mathbf{x}_{i,h}^{l+1} = \mathbf{x}_{i,h}^{l} + \frac{\sum_{j=1}^{n} \mathbf{S}_{ijh}}{\sqrt{n}},$$

while the initial $\mathbf{x}_h$ for each head is just a copy of the input coordinates $\mathbf{x}$. The process is repeated at each layer, where a new normalized displacement matrix is calculated given the updated coordinates, and new SE(3)-equivariant displacement tensor is generated by dot-producting the displacement matrix $\mathbb{D}^l \in \mathbb{R}^{n \times n \times h \times 3}$ and pair-representation $\mathbf{P}^l$ (tensor product is only used in the first layer where displacement matrix has no head dimension).

##### 1.3 Output layers

Final score prediction is given by a dot-product of the final representation of the SE(3)-invariant branch and the SE(3)-equivariant branch, given by:

$$\mathbf{Z} = \mathbf{Z}_{inv}\mathbf{W}_{inv} \odot \mathbf{Z}_{equ}\mathbf{W}_{equ},$$

where $\mathbf{W}_{inv} \in \mathbb{R}^{hd\times o}$ and $\mathbf{W}_{equ} \in \mathbb{R}^{3h\times o}$ are the projection matrices that project the input to desired output dimension o, $\mathbf{Z}_{inv} \in \mathbb{R}^{n\times h\times d}$ is the output of final attention layer following a MLP, and $\mathbf{Z}_{equ} \in \mathbb{R}^{n\times h\times 3}$ is given by:

$$\mathbf{Z}_{equ} = \frac{\sum_{j=1}^{n} \mathbf{S}_{ijh}}{\sqrt{n}}.$$

To predict the translation and rotation score, a two-layer MLP with GeLU activation function is used to get $\mathbf{Z}_{inv}$, taking the first representation in the sequence of the attention output, which corresponds to a padded special token ¡s¿, and projecting it to a length-6 vector. As for the atom-wise diffusion score, another two-layer MLP with GeLU activation function is used to get $\mathbf{Z}_{inv}$, taking the full sequence of the attention layer output, resulting the perturbation score prediction $\mathbf{Z} \in \mathbb{R}^{(n_l+n_p)\times 3}$.

#### 2 Proofs

##### 2.1 Proof for distance matrix being $SE(3)$-invariant

**Lemma 1.** *Let $\mathbf{x} \in \mathbb{R}^{3n}$ be the position coordinates of all $n$ atoms in a molecule, and let $\mathbf{D} \in \mathbb{R}^{n\times n}$ be the distance matrix such that $\mathbf{D}_{ij} = ||\mathbf{x}_i - \mathbf{x}_j||_2$. For any element $g \in SE(3)$, representing a roto-translational operation in 3D Euclidean space, if we apply $g$ to $x$ to obtain new coordinates $x'$ such that $x'_i = g \cdot x_i$ for any atom $i$, and a new distance matrix $\mathbf{D}'$ such that $\mathbf{D}'_{ij} = ||\mathbf{x}'_i - \mathbf{x}'_j||_2$, then we have $\mathbf{D}' = \mathbf{D}$.*

*Proof.* Since each roto-translational operation $g$ on a 3D coordinate $x_i$ can be represented by an affine transformation $Rx_i + v$, where $R \in SO(3)$ is a rotational matrix and $v \in \mathbb{R}^3$ is a translational vector, we have:

$$\begin{aligned} ||g\cdot\mathbf{x}_i - g\cdot\mathbf{x}_j||_2^2 &= ||R\mathbf{x}_i + v - (R\mathbf{x}_j + v)||_2^2 = ||R(\mathbf{x}_i - \mathbf{x}_j)||_2^2 \\ &= (\mathbf{x}_i - \mathbf{x}_j)^\top R^\top R(\mathbf{x}_i - \mathbf{x}_j) = ||\mathbf{x}_i - \mathbf{x}_j||_2^2. \end{aligned}$$

The last equality holds since for each element $R$ in $SO(3)$, we have $R^\top R = RR^\top = \mathbf{I}$. Therefore, for each $i, j$ pair, we have:

$$\mathbb{D}'_{ij} = ||\mathbf{x}'_i - \mathbf{x}'_j||_2 = ||g \cdot \mathbf{x}_i - g \cdot \mathbf{x}_j||_2 = ||\mathbf{x}_i - \mathbf{x}_j||_2 = \mathbb{D}_{ij}.$$

□

##### 2.2 Proof for coordinate update strategy being $SE(3)$-equivariant

**Lemma 2.** *Let $\mathbb{D} \in \mathbb{R}^{n\times n\times 3}$ be the normalized displacement tensor such that $\mathbb{D}_{ij} = \frac{\mathbf{x}_i - \mathbf{x}_j}{\|\mathbf{x}_i - \mathbf{x}_j\|_2}$. For any element $g \in SE(3)$, if we apply $g$ to $x$ to obtain new coordinates $x'$, for any transformation $\Psi(\mathbf{x}) = \mathbf{x} + \sum_j \Phi(\mathbf{x})_{ij}\mathbb{D}_{ij}$ with $\Phi((x))$ being $SE(3)$-invariant transformation, the following hold $\Psi(\alpha_g(\mathbf{x})) = \alpha_g(\Psi(\mathbf{x}))$, i.e. $\Psi(\mathbf{x})$ is a $SE(3)$-equivariant transformation.*

*Proof.* As $\alpha_g(\mathbf{x}_i) = R\mathbf{x}_i + v$, we have:

$$\begin{aligned}
\Psi(\alpha_g(\mathbf{x})) &= \alpha_g(\mathbf{x}) + \sum_j \Phi(\alpha_g(\mathbf{x}))_{ij} \frac{\alpha_g(\mathbf{x}_i) - \alpha_g(\mathbf{x}_j)}{\|\alpha_g(\mathbf{x}_i) - \alpha_g(\mathbf{x}_j)\|_2} \\
&= R\mathbf{x} + v + \sum_j \Phi(\mathbf{x})_{ij} \frac{R\mathbf{x}_i + v - R\mathbf{x}_j - v}{\|R\mathbf{x}_i - R\mathbf{x}_j\|_2} \\
&= R\mathbf{x} + v + \sum_j \Phi(\mathbf{x})_{ij} \frac{R(\mathbf{x}_i - \mathbf{x}_j)}{\|\mathbf{x}_i - \mathbf{x}_j\|_2} \\
&= R(\mathbf{x} + \sum_j \Phi(\mathbf{x})_{ij}\mathbb{D}_{ij}) + v = \alpha_g(\Psi(\mathbf{x})).
\end{aligned}$$

□

##### 2.3 Proof of group action under expectation

Let $\mathbf{x} = \{\mathbf{x}_1, \mathbf{x}_2, \ldots, \mathbf{x}_n\}$ be the position coordinates of all $n$ atoms. We can apply the following operations during the diffusion process:

- System Rotation around the center $\bar{\mathbf{x}}$

$$\alpha^{\text{rot}}(\mathbf{R}, \mathbf{x})_i = \mathbf{R}(\mathbf{x}_i - \bar{\mathbf{x}}) + \bar{\mathbf{x}},$$

  where $\mathbf{R} \in SO(3)$ is a rotation matrix.

- System Translation

$$\alpha^{\text{tr}}(\mathbf{r}, \mathbf{x})_i = \mathbf{x}_i + \mathbf{r},$$

where $\mathbf{r} \in \mathbb{R}^3$ is a translation vector.

– Perturbation Noise

$$\mathbf{x}'_i = \mathbf{x}_i + \boldsymbol{\epsilon}_i,$$

where $\boldsymbol{\epsilon}_i \sim \mathcal{N}(\mathbf{0}, \sigma^2\mathbf{I})$ are independent zero-mean Gaussian noises.

To prove the operations forms a group action, we need to prove that the combined operation $\alpha$, defined as:

$$\alpha(g, \mathbf{x}')_i = \mathbf{R}(\mathbf{x}'_i - \bar{\mathbf{x}}') + \bar{\mathbf{x}}' + \mathbf{r},$$

with $g = (\mathbf{R}, \mathbf{r})$, forms a group action on the noised point cloud $\mathbf{x}'$.

*Proof.* First, note that the noise has zero mean:

$$\mathbb{E}[\boldsymbol{\epsilon}_i] = \mathbf{0} \implies \mathbb{E}[\mathbf{x}'_i] = \mathbf{x}_i.$$

The expected center of the perturbed point cloud keeps the same as original point cloud:

$$\mathbb{E}[\bar{\mathbf{x}}'] = \mathbb{E}\left[\frac{1}{N}\sum_{i=1}^{N}\mathbf{x}'_i\right] = \bar{\mathbf{x}}.$$

To prove $\alpha$ forms a group action under expectation, we need to prove the identity and the compatibility property of $/alpha$.

**Proof of identity**: Let $e = (\mathbf{I}, \mathbf{0})$ be the identity element. Then:

$$\alpha(e, \mathbf{x}')_i = \mathbf{I}(\mathbf{x}'_i - \bar{\mathbf{x}}') + \bar{\mathbf{x}}' + \mathbf{0} = \mathbf{x}'_i.$$

Taking expectation:

$$\mathbb{E}[\alpha(e, \mathbf{x}')_i] = \mathbb{E}[\mathbf{x}'_i] = \mathbf{x}_i.$$

Thus, the identity property holds in expectation.

**Proof for Compatibility Property**:

Let $g_1 = (\mathbf{R}_1, \mathbf{r}_1)$ and $g_2 = (\mathbf{R}_2, \mathbf{r}_2)$. Their composition is:

$$g_1 g_2 = (\mathbf{R}_1\mathbf{R}_2, \mathbf{r}_2 + \mathbf{r}_1).$$

We need to show:

$$\mathbb{E}[\alpha(g_1 g_2, \mathbf{x}')]_i = \mathbb{E}[\alpha(g_1, \alpha(g_2, \mathbf{x}'))_i].$$

Compute the left-hand side (LHS):

$$\begin{aligned}\mathbb{E}[\alpha(g_1g_2,\mathbf{x}')]_i =&\mathbb{E}[\mathbf{R}_1\mathbf{R}_2(\mathbf{x}'_i-\bar{\mathbf{x}}')+\bar{\mathbf{x}}']+\mathbf{r}_2+\mathbf{r}_1\\ =&\mathbf{R}_1\mathbf{R}_2(\mathbf{x}_i-\bar{\mathbf{x}})+\bar{\mathbf{x}}+\mathbf{r}_2+\mathbf{r}_1\end{aligned}$$

Compute the right-hand side (RHS):

First, compute $\mathbb{E}[\alpha(g_2,\mathbf{x}')]_i$:

$$\mathbb{E}[\alpha(g_2,\mathbf{x}')]_i=\mathbf{R}_2(\mathbf{x}_i-\bar{\mathbf{x}})+\bar{\mathbf{x}}+\mathbf{r}_2.$$

The expected center after applying $g_2$ is:

$$\mathbb{E}[\overline{\alpha(g_2,\mathbf{x}')}]=\bar{\mathbf{x}}+\mathbf{r}_2.$$

Then:

$$\mathbb{E}[\alpha(g_2,\mathbf{x}')]_i-\mathbb{E}[\overline{\alpha(g_2,\mathbf{x}')}]=\mathbf{R}_2(\mathbf{x}_i-\bar{\mathbf{x}}).$$

Now, compute the RHS:

$$\begin{aligned}&\mathbb{E}[\alpha(g_1,\alpha(g_2,\mathbf{x}'))_i]\\ &=\mathbb{E}[\mathbf{R}_1[\alpha(g_2,\mathbf{x}')_i-\overline{\alpha(g_2,\mathbf{x}')}]+\overline{\alpha(g_2,\mathbf{x}')}+\mathbf{r}_1]\\ &=\mathbf{R}_1[\mathbb{E}[\alpha(g_2,\mathbf{x}')]_i-\mathbb{E}[\overline{\alpha(g_2,\mathbf{x}')}]]+\mathbb{E}[\overline{\alpha(g_2,\mathbf{x}')}]+\mathbf{r}_1\\ &=\mathbf{R}_1\mathbf{R}_2(\mathbf{x}_i-\bar{\mathbf{x}})+\bar{\mathbf{x}}+\mathbf{r}_2+\mathbf{r}_1\end{aligned}$$

Comparing LHS and RHS:

$$\mathbb{E}[\alpha(g_1g_2,\mathbf{x}')]_i=\mathbb{E}[\alpha(g_1,\alpha(g_2,\mathbf{x}'))_i].$$

Thus, the compatibility property holds in expectation. □

##### 2.4 Diffusion form of SMLD and DDPM

Below we provide a straight-forward derivation for SMLD and DDPM diffusion process. For forward diffusion process in SMLD with Gaussian kernel $P(\mathbf{x}_i|\mathbf{x}_{i-1}) = \mathcal{N}(\mathbf{x}_i; \mathbf{x}_{i-1}, (\sigma_i^2 - \sigma_{i-1}^2)\mathbf{I})$

$$\begin{aligned}
x_i &= x_{i-1} + \sqrt{\sigma_i^2 - \sigma_{i-1}^2}\epsilon \\
&= x_{i-2} + \sqrt{\sigma_{i-1}^2 - \sigma_{i-2}^2}\epsilon' + \sqrt{\sigma_i^2 - \sigma_{i-1}^2}\epsilon \\
&= x_{i-2} + \sqrt{\sigma_i^2 - \sigma_{i-2}^2}\epsilon \\
&= \cdots \\
&= x_0 + \sqrt{\sigma_i^2 - \sigma_0^2}\epsilon \\
&= x_0 + \sigma_i\epsilon
\end{aligned}$$

If we assume $\sigma_0 = 0$, and notice for two independent gaussian variable $\epsilon_1 \sim \mathcal{N}(0, \sigma_1^2)$ and $\epsilon_2 \sim \mathcal{N}(0, \sigma_2^2)$, $\epsilon_1 + \epsilon_2 \sim \mathcal{N}(0, \sigma_1^2 + \sigma_2^2)$.

Similarly, for forward diffusion process in DDPM with Gaussian kernel $P(\mathbf{x}_i|\mathbf{x}_{i-1}) = \mathcal{N}(\mathbf{x}_i; \sqrt{1-\beta_i}\mathbf{x}_{i-1}, \beta_i\mathbf{I})$

$$\begin{aligned}
x_i &= \sqrt{1-\beta_i}x_{i-1} + \sqrt{\beta_i}\epsilon;\ \epsilon \sim \mathcal{N}(0,1) \\
&= \sqrt{1-\beta_i}(\sqrt{1-\beta_{i-1}}x_{i-2} + \sqrt{\beta_{i-1}}\epsilon') + \sqrt{\beta_i}\epsilon \\
&= \sqrt{(1-\beta_i)(1-\beta_{i-1})}x_{i-2} + \sqrt{(1-\beta_i)\beta_{i-1}}\epsilon' + \sqrt{\beta_i}\epsilon \\
&= \sqrt{(1-\beta_i)(1-\beta_{i-1})}x_{i-2} + \sqrt{(1-\beta_i)\beta_{i-1} + \beta_i}\epsilon \\
&= \sqrt{(1-\beta_i)(1-\beta_{i-1})}x_{i-2} + \sqrt{1-(1-\beta_i)(1-\beta_{i-1})}\epsilon \\
&= ... \\
&= \sqrt{\bar{\alpha}_t}x_0 + \sqrt{1-\bar{\alpha}_i}\epsilon;\ \bar{\alpha}_i = \prod_{s=0}^{i}(1-\beta_s)
\end{aligned}$$

### 3 Evaluation Details

We compared DTMol with two search-based docking methods Smina and gnina, and with two previously reported high-accurate machine-learning based docking methods Unimol and DiffDock.

*SMINA* Koes et al. (2013) uses Autodock-Vina docking kernel with a new scoring function. We set '–exhaustiveness' to 100 (default is 8) for better performance, and set '–num_modes' to generate 10 candidates. Other settings are kept as default.

*GNINA* McNutt et al. (2021) equipped with an additional 3D CNN for scoring function. Same to Smina, we set the '–exhaustiveness 100' and '–num_modes' to 10.

*Unimol* Zhou et al. (2022) is a universal molecular embedding model for multiple downstream tasks, including molecular docking. We directly use the weights pretrained on binding pose prediction provided by Unimol, and all parameters are kept as default. For a fair comparison, all pockets are the same according to different pocket-based docking models.

*DiffDock* Corso et al. (2023) is a generative diffusion model designed for blind docking. As it is not able to specify a protein pocket with DiffDock, we generated 50 samples on the whole protein with DiffDock, and then selected the 10 samples with minimum RMSD as the final collection. We calculate top1 and top5 scores among them according to the score given by the confidence model.

*DiffBindFR* Zhu et al. (2024) is another generative diffusion model designed for pocket-based molecular docking. Similar to DiffDock, it employs a graph neural network with a torsion diffusion process. Additionally, the model considers protein side chain flexibility and allows specifying a protein pocket with a reference ligand as the starting position for the reverse diffusion process (docking). We used the default settings of the model, generating 40 samples for each protein-ligand pair, and used the Smina scoring function to rank the predictions.

*DTMol* run with Smina as a scoring function. Multiple samples (default is 10) are generated by DTMol for each docking process, which are then scored and ranked by Smina.

| Parameter | Value |
|---|---|
| Learning Rate | **0.001**, 0.0001, 0.00001 |
| Start Learning Rate Factor | **0.001**, 0.01, 0.1 |
| Learning Scheduler | **Linear**, CosineAnnealing, CosineAnnealingWarmStart |
| Maximum Scale of Time Embedding | $\mathbf{T^{1.5}}$, $2T$ |
| $\sigma^{tr}_{\max}$ | **4**, 8, 12 |
| $\sigma^{pert}_{\max}$ | **2**, 3, 4 |
| Shared attention between branches | **Yes**, No |
| Form for translation and rotation noise | **VE**, VP |
| Form for perturbation noise | VE, **VP** |
| Max reverse diffusion time | **20**, 30, 40 |

**Table S1.** Hyperparameters searched for the DTMol model; the final parameters are highlighted in bold.

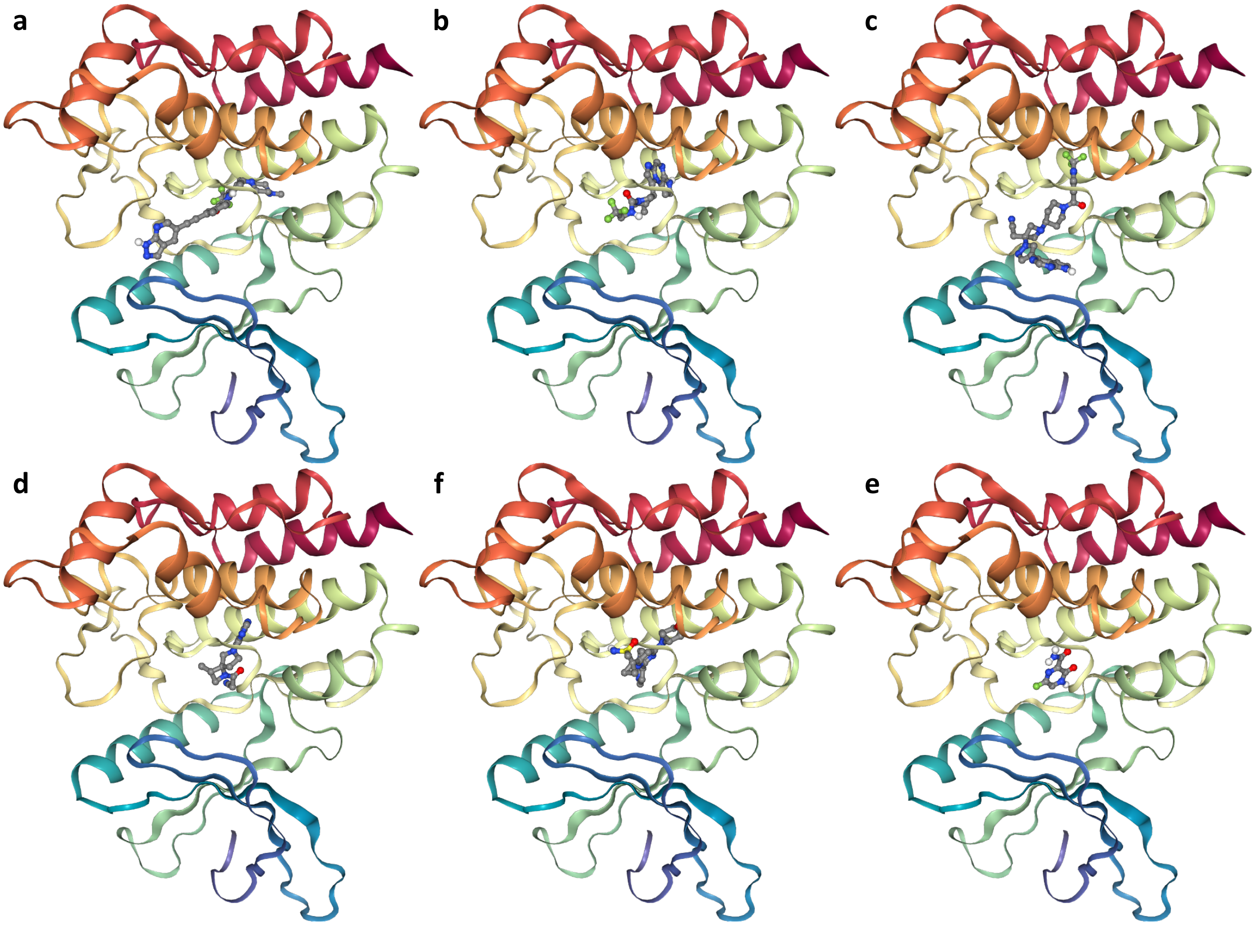


**Supplementary Figure S1.** An illustration of docked poses generated by DTMol for six drugs binding to the JAK2 protein (PDB ID: 6bbv): (a) HQP1351, (b) Upadacitinib, (c) Itacitinib, (d) Delgocitinib, (e) Ceralascrtib, and (f) Favipiravi.

| Code | Compounds | Experiment Rank | DockRank | DTMol Rank | MD Rank | DiffbindFR Rank |
|---|---|---|---|---|---|---|
| jak11 | Upadacitinib | 1 | 18 | 4 | 12 | 21 |
| jak1 | HQP1351 | 2 | 2 | 16 | 1 | 5 |
| jak27 | Itacitinib | 3 | 5 | 5 | 11 | 24 |
| jak16 | Delgocitinib | 4 | 24 | 6 | 21 | 27 |
| jak30 | Ceralasertib | 5 | 19 | 8 | 22 | 26 |
| jak29 | Favipiravir | 6 | 26 | 7 | 27 | 19 |
| jak14 | DB04716 | 7 | 21 | 14 | 18 | 3 |
| jak26 | Regadenoson | 8 | 9 | 26 | 23 | 25 |
| jak19 | DB07794 | 9 | 17 | 21 | 20 | 6 |
| jak32 | BMS-911543 | 10 | 8 | 12 | 10 | 7 |
| jak7 | Cerdulatinib | 11 | 11 | 13 | 7 | 8 |
| jak12 | A-674563 | 12 | 20 | 27 | 26 | 10 |
| jak18 | Cetrimonium | 13 | 27 | 23 | 6 | 23 |
| jak17 | Fenebrutinib | 14 | 1 | 20 | 2 | 13 |
| jak20 | IMR-687 | 15 | 14 | 1 | 16 | 20 |
| jak2 | Lorecivivint | 16 | 3 | 22 | 8 | 4 |
| jak25 | PF-06651600 | 17 | 25 | 9 | 19 | 22 |
| jak23 | PP-121 | 18 | 22 | 15 | 25 | 9 |
| jak22 | Osoresnontrine | 19 | 23 | 3 | 24 | 2 |
| jak3 | Entrectinib | 20 | 4 | 17 | 5 | 14 |
| jak5 | Gandotinib | 21 | 7 | 18 | 3 | 17 |
| jak10 | CC-401 | 22 | 15 | 24 | 17 | 12 |
| jak8 | DB08073 | 23 | 12 | 19 | 13 | 16 |
| jak21 | PF-04447943 | 24 | 16 | 2 | 15 | 1 |
| jak4 | Filgotinib | 25 | 6 | 11 | 14 | 15 |
| jak9 | CHIR-124 | 26 | 13 | 10 | 9 | 18 |
| jak6 | Capivasertib | 27 | 10 | 25 | 4 | 11 |

**Table S2.** Comparison of the ranks given by the assay experiment (Experiment Rank), docking (DockRank), DTMol (AI Rank), and docking followed by MD relaxation (MD Rank) for different compounds. DTMol got the only positive Spearman's rank correlation (0.192), while the DiffbindFR, the conventional docking method and conventional docking followed by molecular dynamic relaxation got negative correlation scores (-0.264, -0.146 and -0.194).
